# Systematic Analysis of Human Tissue- and Cell-Specific Metabolic Models Identifies High-Sugar, High-Fat Diet–Induced Liver Dysregulation

**DOI:** 10.64898/2026.02.18.706568

**Authors:** Mengzhen Li, Mengnan Shi, Cheng Zhang, Hasan Turkez, Mathias Uhlen, Adil Mardinoglu

## Abstract

Human tissues exhibit specialized metabolic functions that are essential for maintaining whole-body metabolic homeostasis. To systematically characterize organ- and cell-type-specific metabolic heterogeneity, we constructed 32 tissue-specific and 81 cell-type-specific enzyme-constrained genome-scale metabolic models (ecGEMs) by integrating the global human metabolic network with the tissue- and single-cell transcriptomic data from the Human Protein Atlas (HPA). Our analysis revealed pronounced differences in metabolic network architecture and activity across the human body, identifying key cell types that drive tissue metabolic functions. To demonstrate the applicability of these models, we employed the liver-specific ecGEM to investigate the metabolic reprogramming induced by a high-sugar, high-fat (HSHF) diet, a primary driver of metabolic dysfunction-associated fatty liver disease (MAFLD). Flux balance analysis revealed a fundamental transition in hepatic metabolism: from a flexible, multi-functional system toward a constrained, lipid-centric regime. This state is characterized by carbohydrate and lipid overload, mitochondrial respiratory dysfunction, and a compromised capacity for reactive oxygen species (ROS) detoxification. These computational predictions were validated through integrative analysis of transcriptomic data from a human MAFLD cohort and metabolomic profiles from an *in vivo* HSHF rat model. Together, this work provides a comprehensive atlas of human metabolic models, enabling the systematic investigation of metabolic features across tissues and conditions from a systems-level perspective.

**Graphical Abstract:** 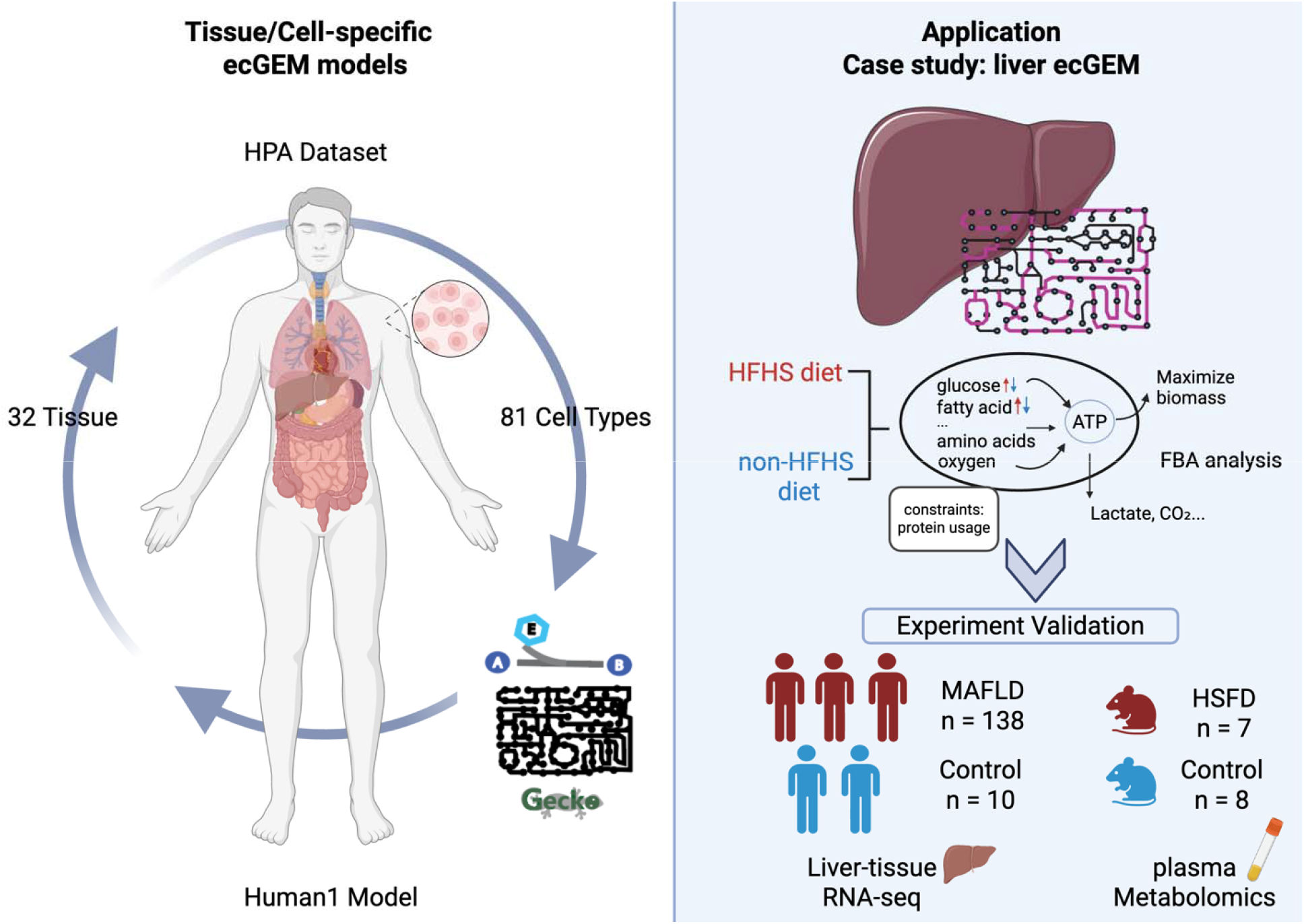

## Introduction

Understanding human metabolism is fundamental for advancing our knowledge of both physiological homeostasis and disease pathophysiology^1^. The human body is a complex system in which distinct tissues and cell types exhibit specialized metabolic programs closely linked to their functional and pathological roles. The progression of systemic metabolic disorders, including obesity, diabetes, cardiovascular disease, and metabolic dysfunction-associated fatty liver disease (MAFLD), is invariably driven by the reprogramming of tissue-specific metabolic functions ^2^. Consequently, a high-resolution characterization of tissue- and cell-specific metabolic profiles, particularly their change under dietary or pathological stress, is essential for elucidating the disease mechanisms and identifying precision therapeutic biomarkers.

To study these complex systems, Genome-scale metabolic models (GEMs) have emerged as a powerful tool in systems biology. By translating the metabolite and reaction networks into mathematical frameworks, GEMs enable the quantitative characterization of metabolic states under specific conditions ^3,4^. Among the reconstructed GEMs, Human1 and Recon3 are comprehensive global human GEMs that integrate reactions, metabolites, and associated genes across the human body. In Human1, genes encode enzymes, and Gene–Protein–Reaction (GPR) rules are used to link genes to the metabolic reactions they catalyze ^6^. Despite the availability of well-established global human GEMs and the generation of numerous tissue- and cell-specific models, the metabolic features of many individual tissues and cell types remain incompletely characterized. By incorporating tissue- and cell-specific gene expression patterns, Human1 can be extended to infer metabolic networks and flux distributions at both the tissue and cellular levels.

The Human Protein Atlas (HPA) is a comprehensive resource that maps human genes and proteins across tissues and cells. It includes bulk transcriptomics data for 32 human tissues^7^ and single-cell transcriptomics data for 81 distinct cell types^8,9^. In this study, we constructed tissue- and cell-specific GEMs by integrating transcriptomics data from the HPA with the Human1 global model. Our analysis systematically revealed the tissue- and cell-specific metabolic profiles and characterized their distinctive features. To demonstrate the applicability of these models, we analyzed liver enzyme-constrained genome-scale metabolic models (ecGEMs) under different dietary conditions. We finally validated the predicted metabolic alterations using the transcriptomic data of MAFLD patients and metabolomic data from diet-specific *in vivo* rat models.

## Results

### Generation of Tissue-Specific GEMs to Characterize Metabolic Functions

First, to capture tissue-specific metabolic features, 2,897 protein-coding genes from HPA were mapped into Human1. We found that most of these protein-coding genes showed low tissue specificity (Figure 1A). This suggests that the associated metabolic reactions constitute a “core” metabolic network, reflecting fundamental biological processes that are essential for cellular maintenance and survival across diverse tissue types. Moreover, around 10% of these genes were tissue-enriched, particularly in the liver, suggesting that specific metabolic activities are more prominent in certain tissues.

**Fig. 1:**
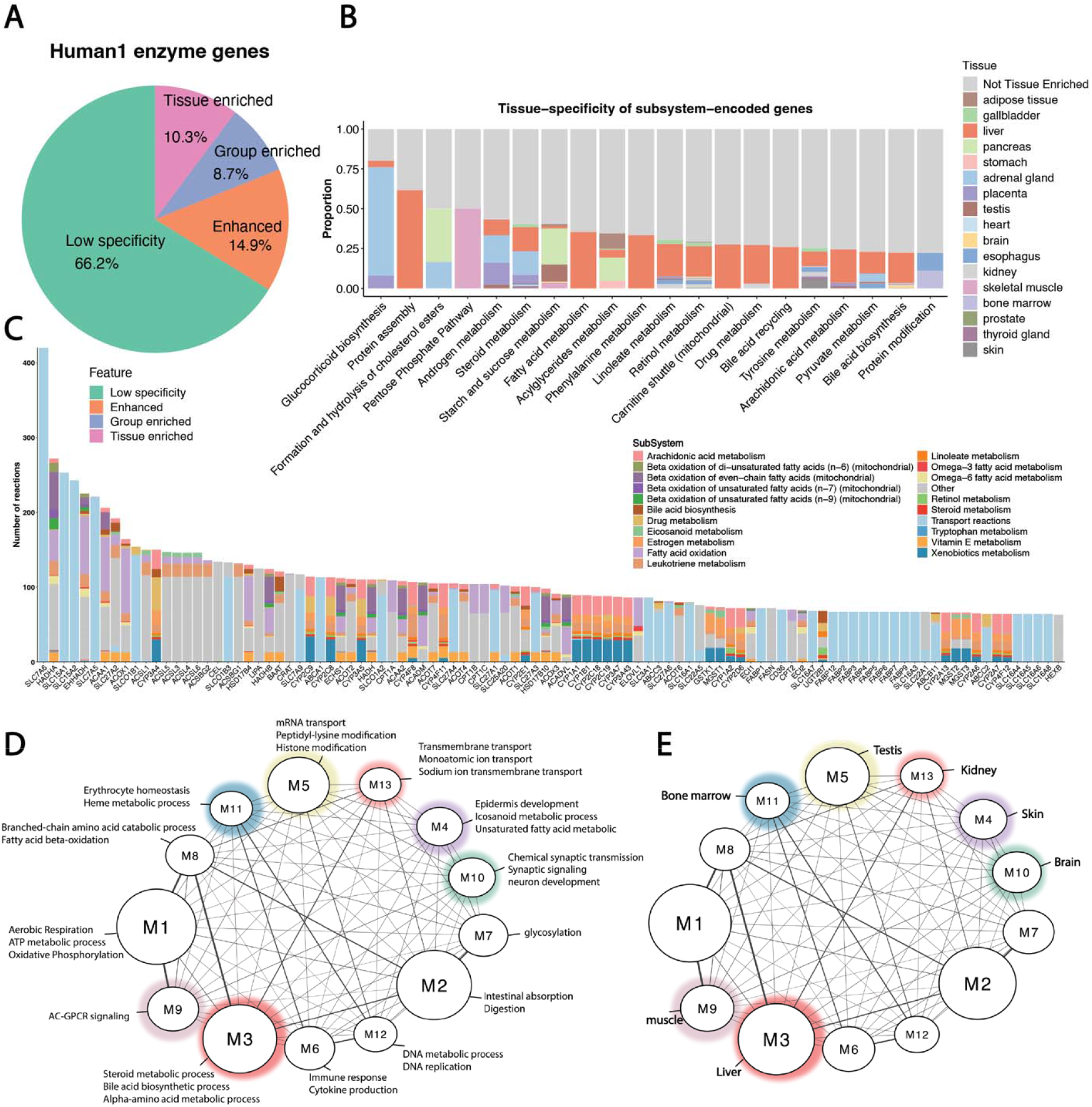
Tissue-specific characteristics of enzyme-encoding genes in the Human1 model. (A) Pie chart showing the proportion of tissue-specific genes among 2897 enzyme-encoding genes in Human1, categorized as tissue-enriched, group-enriched, enhanced, or low-specificity. (B) Top 20 metabolic subsystems whose enzyme-encoding genes display high tissue specificity, indicating that reactions within these subsystems are more tissue-restricted. (C) Top 100 enzyme-encoding genes catalyzing multiple metabolic reactions, grouped by their associated subsystems. (D) Weighted gene co-expression network analysis (WGCNA) of 2,897 enzyme-encoding genes identified 13 co-expression modules with distinct metabolic functions. (E) Correlation analysis between module eigengenes and tissue types. Module eigengenes, representing the first principal component of each co-expression module, were correlated with tissue indicator variables using Pearson correlation. Modules showing strong correlations (r > 0.7) were considered tissue-specific, highlighting distinct metabolic functions across tissues.

The 2,897 protein-coding genes participate in 13,073 reactions, which are further categorized into 150 different subsystems in the Human1 model. We next identified the top subsystems whose genes exhibited a high proportion of tissue-enriched expression. As shown in Figure 1B, most of the genes responsible for glucocorticoid biosynthesis were enriched in the adrenal gland, while a large proportion of pentose phosphate pathway enzyme genes were enriched in skeletal muscle. In addition, liver shows up as a central and unique tissue in these metabolic pathways compared to other tissues, that genes in several subsystems, such as protein assembly, fatty acid metabolism, and phenylalanine metabolism, were predominantly expressed in the liver. We also identified the top genes associated with multiple metabolic reactions, highlighting their essential roles in human metabolism (Figure 1C). For instance, members of the SLC family were primarily associated with transport reactions, whereas HADHA, EHHADH, and ACAA1 were involved in several subsystems, including fatty acid oxidation and leukotriene metabolism.

Next, we aimed to investigate the 2,897 protein-coding genes at the gene module level. We obtained RNA-seq data from 200 samples across 32 different tissues from the HPA. Using Weighted Gene Co-expression Network Analysis (WGCNA), we identified 13 distinct gene modules from these genes, with module sizes ranging from 49 to 448 genes (Figure 1D, Supplementary Table1). Each module exhibited significant enrichment for specific Gene Ontology (GO) pathways. As expected, many of these modules were functionally related to metabolic activity. For example, Module 1 was associated with essential energy production processes such as aerobic respiration, ATP generation, and oxidative phosphorylation (OXPHOS). Module 3 was enriched in steroid metabolism and bile acid biosynthesis, closely reflecting liver-specific functions. Interestingly, correlation analysis between module-specific genes and tissue indicator variables (Figure 1E) showed that Module 3 was highly correlated with liver tissue (correlation score = 0.93). Similarly, Module 11, enriched in heme metabolism, was strongly correlated with bone marrow (correlation score = 0.97), while Module 4, associated with eicosanoid metabolism, showed a strong correlation with skin (correlation score = 0.79). In contrast, Module 1, which covers fundamental energy metabolism, did not exhibit clear tissue specificity, suggesting that these metabolic reactions are ubiquitous across tissues.

### Metabolic Features Correlating Between Tissues and Their Constituent Cell Types

In order to generate tissue- and cell-specific ecGEMs, RNA-seq data from 32 tissues and single-cell data from 81 cell types were obtained from the HPA. Mean expression values were calculated across tissues and cell types to generate representative ecGEMs, using Human1 as the reference model. To explore structural differences between tissues and cells, we applied t-distributed Stochastic Neighbor Embedding (t-SNE) analysis using binary presence/absence data for 13,073 reactions across tissue- and cell-specific ecGEMs. As shown in Figure 2A, kidney tissue and three cell types, including Paneth cells, inhibitory neurons, and exocrine glandular cells, were outliers, indicating distinct metabolic characteristics compared to others. When focusing on the central overlapping region, we observed that tissue samples were positioned near their corresponding cell types, suggesting similar metabolic structures between tissues and their corresponding cells. To further elucidate the interplay between tissue-level and cellular metabolic features, we utilized the COMPASS algorithm^10^, leveraging the Recon3 Global human GEM, to infer metabolic activity profiles across various tissues and cell types. This approach enabled the quantification of 6,135 distinct metabolic reactions across 95 subsystems. Consistent with our previous findings, tissues exhibited higher correlation scores with their corresponding cell types. For each tissue, we identified and highlighted the cell type with the highest correlation score and designated it as the representative cell type for that organ (Figure 2B), for example, hepatocytes in the liver, cardiomyocytes in the heart, proximal tubular cells in the kidney, and alveolar cells in the lung.

**Fig. 2:**
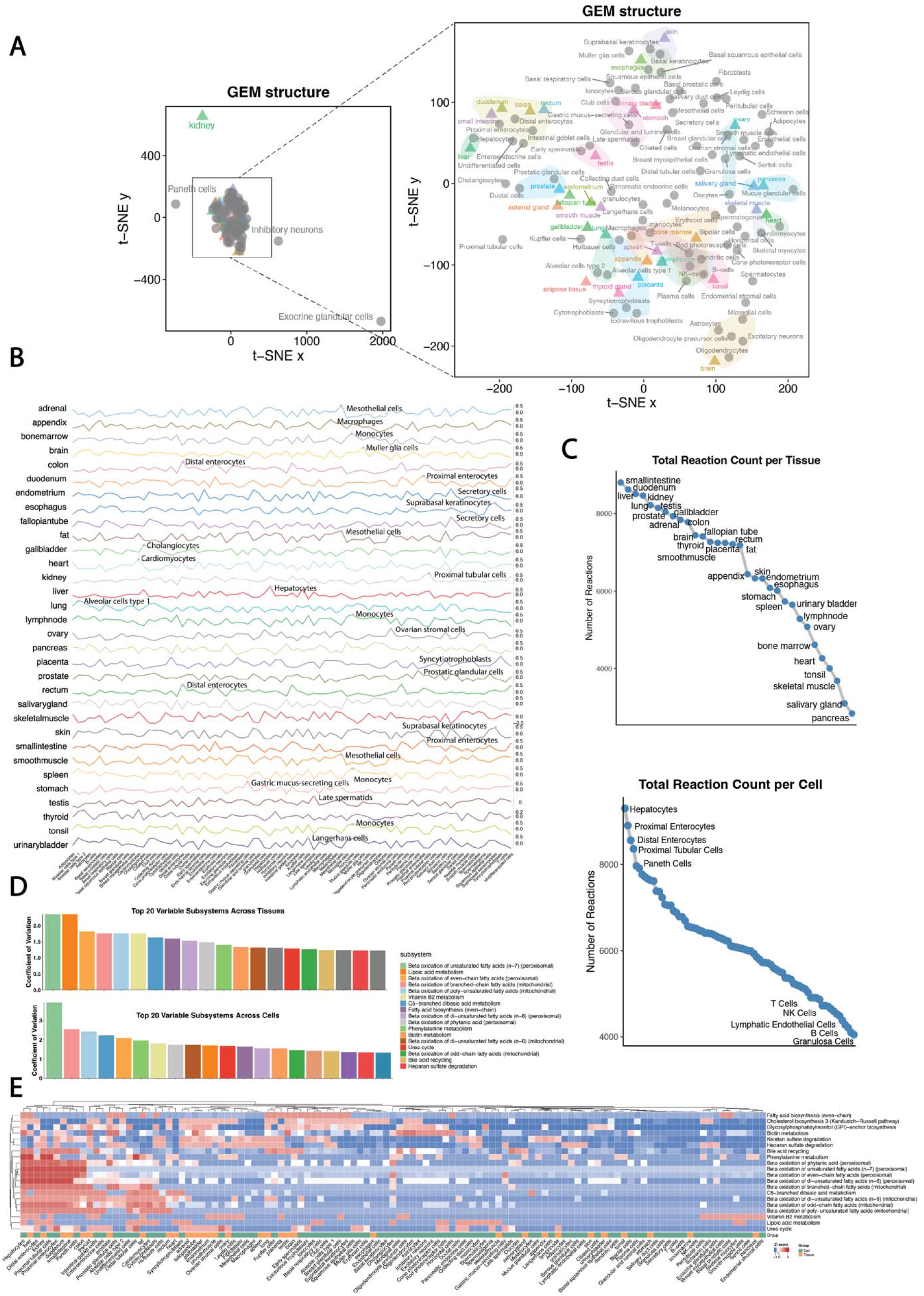
Correlated metabolic features between tissues and corresponding cell types. (A) t-SNE visualization of 32 tissue and 81 cell-type ecGEMs, illustrating structural differences among models. Tissues and their corresponding cell types positioned in close proximity indicate similar metabolic features. (B) Spearman correlations between tissue and cell types based on COMPASS-inferred metabolic reaction activity. The cell type showing the highest correlation with each tissue is labeled on the plot. (C) Total number of active reactions in tissue and cell ecModels, showing distinct metabolic reaction counts. (D) The top 20 most variable metabolic subsystems identified across tissue and cell ecGEMs showed substantial concordance, with 17 subsystems shared between the two rankings. (E) Heatmap showing the z-scores of reactions counts in the selected subsystems across tissue and cell ecModels. Functionally related tissues and cells cluster together based on their metabolic similarity.

Besides, the number of activated reactions varied across tissue- and cell-specific ecGEMs. As shown in Figure 2C, digestive-related tissues, including the small intestine, duodenum, liver, and kidney, exhibited a higher number of active reactions, consistent with their corresponding cell types, such as hepatocytes, enterocytes, and renal tubular cells. In contrast, immune cells such as T cells, B cells, and NK cells exhibited fewer activated reactions, indicating a resting metabolic state. Among all subsystems, top 20 variable subsystems across tissues and cell types are highly overlapped, with fatty acid–related subsystems exhibiting the highest variance in reaction numbers for both tissue and cell ecGEMs, indicating substantial differences in fatty acid metabolism across the organ (Figure 2D). To further examine the distribution of reactions in these subsystems, we generated a heatmap (Figure 2E), which revealed that functionally related tissues and cells clustered together and displayed distinct patterns.

### Different Metabolic Features Between Different Tissues and Cell Types in the Human Body

After establishing the close structural relationship between tissue- and cell-specific ecGEMs, we next focused on differences in reaction activity as inferred by COMPASS and identified significantly different reactions in each tissue compared to all others. As shown in Figure 3A, the liver exhibited the highest number of upregulated metabolic reactions, followed by the kidney, whereas the pancreas, bone marrow, and skeletal muscle displayed higher numbers of significantly downregulated reactions. We selected two representative tissues, the liver and pancreas, and examined the most significantly different reactions between them and other tissues (Figure 3D). In the liver, multiple isoforms of pyruvate kinase (PK2, PK3, PK4) and 3α-hydroxysteroid dehydrogenase were significantly upregulated. Reactions related to vitamin and amino acid metabolism, including retinol dehydrogenase (all-trans, NADPH) and mitochondrial glutamate dehydrogenase, also showed strong activity, highlighting the liver’s central role in metabolism. In contrast, in the pancreas, enzymes involved in nucleotide metabolism, carbohydrate metabolism, heme biosynthesis, and steroid metabolism were predominantly downregulated, suggesting a relatively reduced metabolic activity in these pathways compared with other tissues.

**Fig. 3:**
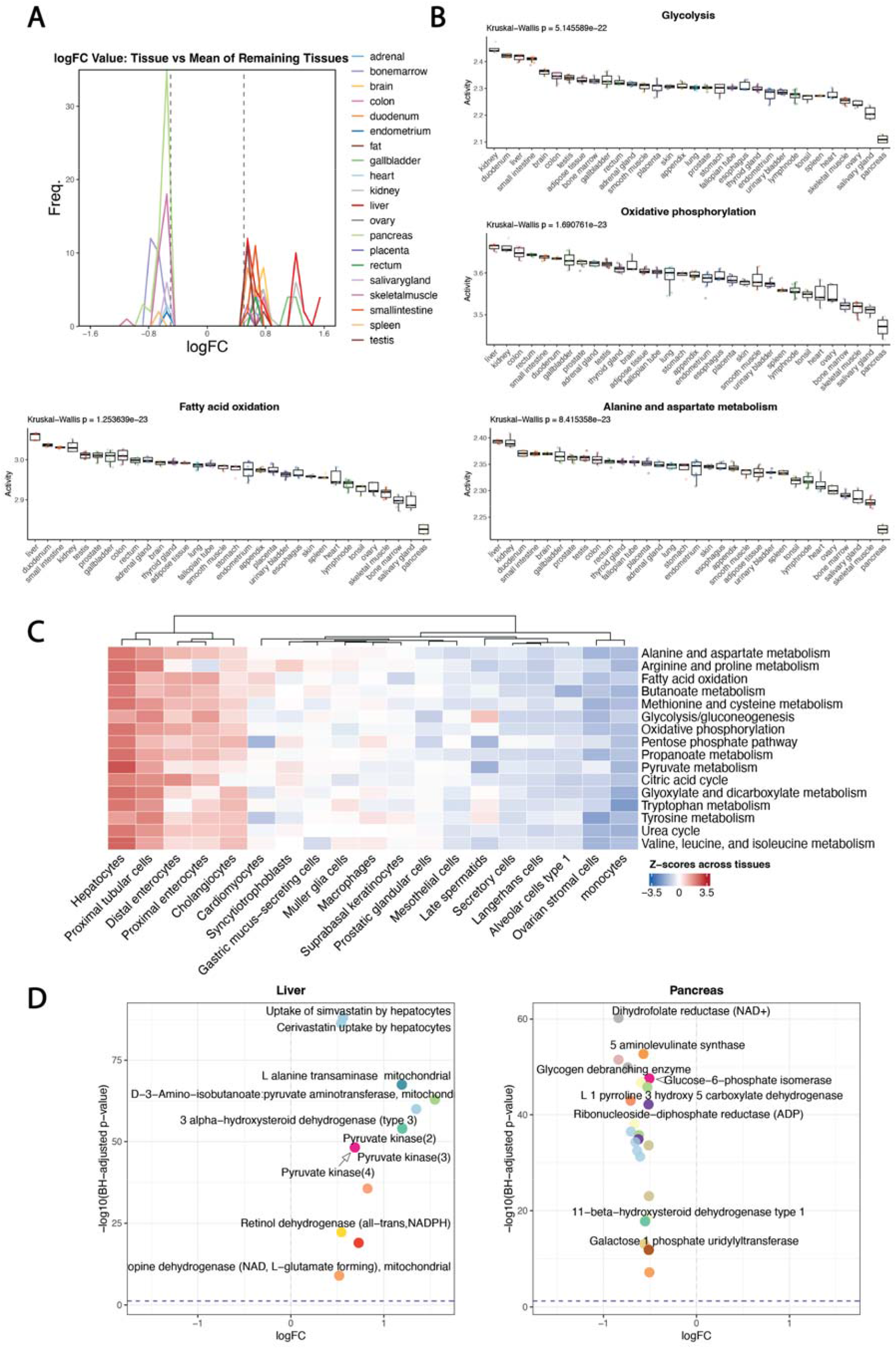
Diversity of metabolic features across human tissues and cell types. (A) Numbers of significantly upregulated or downregulated metabolic reaction activities in each tissue compared with all others. The activity of 61,135 metabolic reactions across tissues was inferred using COMPASS, and statistical significance was assessed using the limma package (B) Mean activity of metabolic reactions in four energy production–related subsystems— glycolysis, oxidative phosphorylation (OXPHOS), fatty acid oxidation, and alanine/aspartate metabolism—showed clear tissue-specific differences, with higher activity observed in the liver, kidney, and digestive-related tissues. (C) Mean metabolic activity of selected energy production pathways across cell types. Hepatocytes and enterocytes exhibited higher activity compared with other cell types. (D) The most significantly altered metabolic reactions in the liver and pancreas relative to other tissues. The liver, a highly metabolically active organ, exhibited increased activity in reactions associated with amino acid metabolism and energy-related pathways. In contrast, the pancreas showed reduced activity in pathways related to nucleotide metabolism, carbohydrate metabolism, and heme biosynthesis.

Focusing on energy production, we selected four subsystems (glycolysis, OXPHOS, fatty acid oxidation, and alanine/aspartate metabolism) to compare differences across tissues (Figure 3B). We found that reactions involved in these subsystems were most strongly activated in the liver, kidney, and digestive tissues of lower gastrointestinal tract, indicating high energy demand in these organs. Similarly, at the cell type level, energy production systems were more active in hepatocytes, tubular cells and enterocytes (Figure 3C).

### Application of the Liver ecGEM Demonstrates Reliability in Capturing Metabolic Features

Next, to demonstrate the application potential of the established ecGEMs, we selected the liver as a representative tissue and performed FBA based on the liver-specific ecGEM. We defined two dietary conditions: a high-fat, high-sugar (HSFD) diet and an HSFD -constrained diet. To mimic dietary changes in the ecGEMs, the uptake rates of glucose and fatty acids were set to their maximum limits under the HFHS condition, while in the HFHS-constrained condition, the uptake rates were restricted. FBA was then performed to calculate the flux distribution of each reaction under both conditions. As shown in Figure 4A, under the HFHS diet condition, fluxes in the mitochondrial carnitine shuttle, steroid metabolism, glycolysis, and the pyruvate system were upregulated compared with the HFHS-constrained diet. We then examined the results at the reaction level. As shown in Figure 4B, the flux through hydrogen-peroxide oxidoreductase was decreased, indicating a reduced antioxidative capacity under the HSFD diet. Additionally, the hydrolysis of cholesteryl-EPA was downregulated, potentially leading to the accumulation of cholesteryl esters in the liver. In contrast, D-glucose 6-phosphotransferase and fatty acid oxidation were upregulated, while several reactions in phospholipid metabolism were downregulated. We further validated these findings in a cohort of MAFLD patients with liver tissue RNA-seq data, observing a similar pattern: oxidative phosphorylation and the respiratory electron transport chain were downregulated, accompanied by upregulation of glycolytic processes through glucose-6-phosphate.

**Fig. 4:**
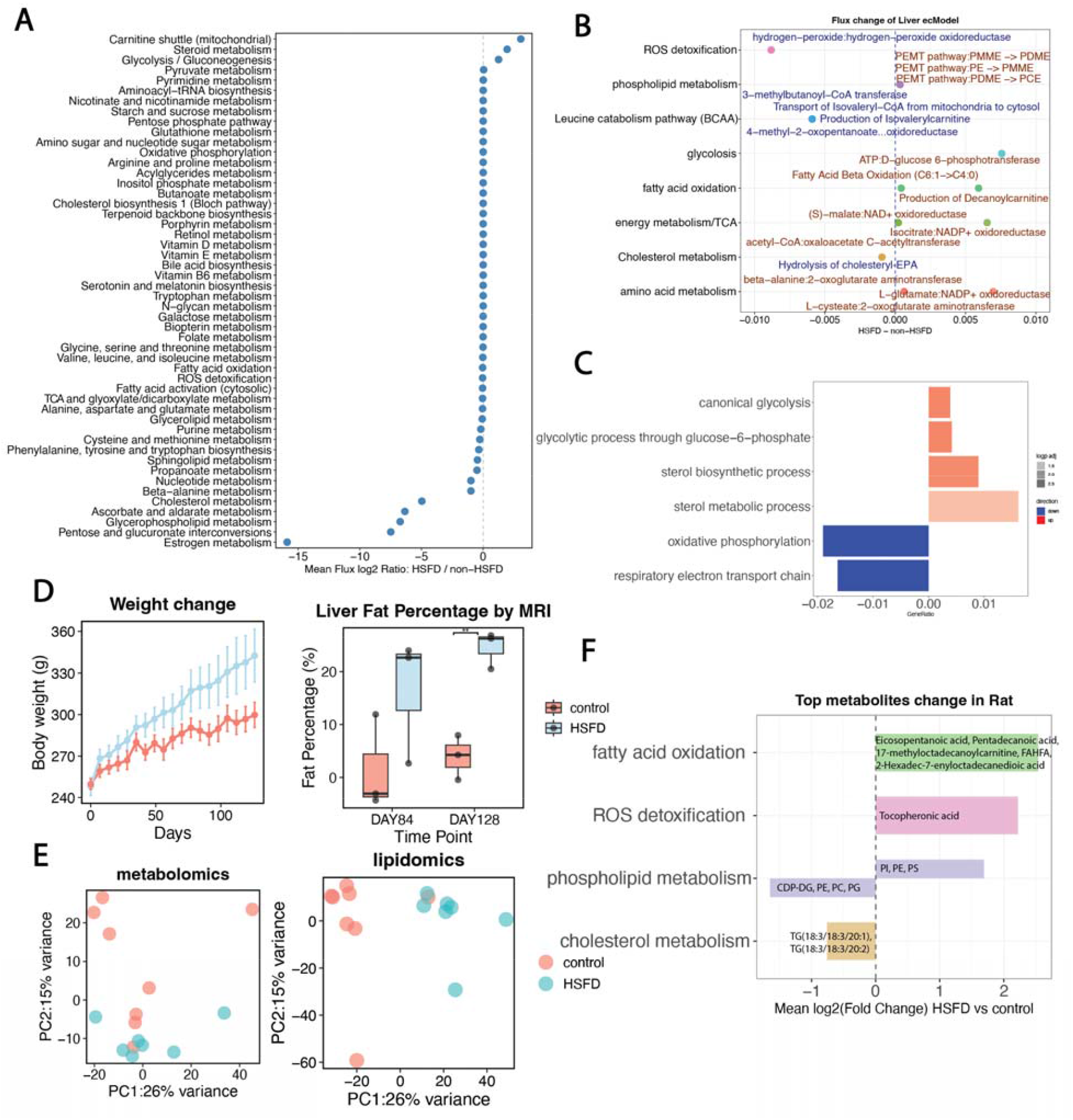
Application of the liver ecGEM: Metabolic alterations under two dietary modes. (A) Subsystem flux changes under two dietary conditions—a high-fat, high-sugar diet (HSFD) and an HSFD-constrained diet—revealed upregulation of the mitochondrial carnitine shuttle, steroid metabolism, glycolysis, and the pyruvate system in the HSFD group. (B) Reaction-level flux differences showed decreased activity of hydrogen-peroxide oxidoreductase, while D-glucose 6-phosphotransferase and fatty acid oxidation were upregulated. (C) Validation using RNA-seq data from MAFLD liver tissues demonstrated a similar trend: oxidative phosphorylation and the respiratory electron transport chain were downregulated, accompanied by upregulation of glycolytic processes via glucose-6-phosphate. (D) *In vivo* validation in a rat model showed that the HSFD group had greater body weight gain over 126 days and developed fatty liver, as confirmed by MRI imaging. (E) PCA analysis of metabolomics and lipidomics data revealed distinct metabolic profiles between diet groups. (F) The most significantly altered metabolites between the two diets were validated in an *in vivo* rat model. Consistent with the liver ecGEM flux analysis, fatty acid–related metabolites were markedly upregulated. In addition, tocopheronic acid levels increased, indicating enhanced oxidative stress under the HSFD condition.

To further validate these findings in an *in vivo* model, we conducted a controlled dietary intervention study in rats. Animals in the control group were maintained on a standard diet, whereas the HSFD group received a calorie-adjusted high-sugar–high-fat diet. After 126 days of treatment, plasma samples were collected for metabolomic analysis. As shown in Figure 4D, rats in the HSFD group exhibited a continuous increase in body weight compared to the control group, and liver fat accumulation was observed by MRI (Supplementary Figure 1-2). PCA analysis revealed distinct plasma metabolic and lipidomic profiles between the two groups (Figure 4E). We then examined the most significantly altered metabolites between the diets (Supplementary Table2). Consistent with the liver ecGEM FBA results, fatty acid–related metabolites were significantly upregulated, and phospholipid-related metabolites were also altered. Besides, tocopheronic acid was increased, suggesting an elevated antioxidant demand. Overall, using liver ecGEM, we predicted metabolic changes in liver tissue following HSFD and observed elevated levels of fatty acid oxidation and glycolysis accompanied by oxidative stress. These findings were further validated in a MAFLD cohort and *in vivo* rat models.

## Discussion

In this study, we characterize tissue- and cell-type–specific metabolic features across the human body by integrating HPA transcriptomics data with the Human1 and Recon3 global human GEMs using genome-scale metabolic modeling and flux balance analysis. This integrative metabolic landscape provides a systems-level view of human metabolism and establishes a framework for analyzing metabolic specialization across cells and tissues. To demonstrate the applicability of the generated models, we imposed diet-specific constraints on liver-specific ecGEMs and inferred corresponding changes in metabolic fluxes. The predicted flux alterations were subsequently evaluated using population-based cohorts and diet-specialized *in vivo* models. Together, these analyses demonstrate that the established tissue- and cell-specific metabolic models capture both baseline metabolic states and condition-dependent metabolic reprogramming, thereby supporting the investigation of disease-associated metabolic alterations from a systems-metabolism perspective.

Previous studies examining organ-specific metabolic features have largely relied on metabolic profiling from clinical samples or animal models. However, these profiling methods primarily capture state-dependent associations rather than functional flux distributions. Without accounting for network-level constraints and dynamic flux changes, it remains challenging to fully understand how metabolic pathways are redistributed under varying physiological conditions. Furthermore, these methods are dependent on the availability of physical samples. In contrast, the ecGEMs developed in this study offer a complementary framework. Through *in silico* simulations, these models enable the prediction of tissue- or cell-specific metabolic responses to novel dietary or pharmacological perturbations. Moreover, by integrating enzyme, metabolite, and reaction information within a unified network structure, ecGEMs facilitate a comprehensive, system-level understanding of metabolic features.

In this study, we constructed ecGEMs from transcriptomic data across multiple human organs and tissues, representing their general, stable metabolic states. We first demonstrated that the established models are consistent with current biological knowledge, thereby supporting their reliability. In addition, the application of two independent approaches, including GEM construction and COMPASS-based metabolic activity inference, yielded concordant results, providing further validation of the robustness of our methodological framework. For instance, among all tissues examined, the duodenum and small intestine exhibited the highest metabolic capacity, supported by both the large number of active reactions in tissue-specific ecGEMs and the elevated metabolic activity inferred by COMPASS. This is consistent with their known biological roles, including rapid epithelial turnover, energy-intensive nutrient absorption, and the presence of densely populated immune cell compartments^20,21^. Moreover, the liver plays a central role in systemic metabolism, particularly in the regulation of lipid, glucose, and amino acid metabolism ^22^, as reflected in our findings.

Beyond representing basal metabolic states, the established ecGEMs are also capable of capturing condition-dependent metabolic flux changes. In this study, we applied liver-specific ecGEMs to investigate metabolic alterations induced by an HSHF diet, mimicking MAFLD-like conditions. The predicted flux changes showed strong consistency with observations from disease cohorts and HSHF diet animal models at both the transcriptomic and metabolomic levels. Several key findings emerged from this analysis. First, under HSHF conditions, the liver exhibits carbohydrate and lipid metabolic overload accompanied by mitochondrial dysfunction. The ecModel predicted significantly increased fluxes through glycolysis (e.g., ATP:D-glucose 6-phosphotransferase) and fatty acid β-oxidation (FAO). Consistently, human cohort data revealed significant enrichment of upregulated canonical glycolysis pathways. In contrast, oxidative phosphorylation and the respiratory electron transport chain were downregulated across datasets, suggesting that HSHF-induced nutrient overload may progressively compromise mitochondrial respiratory capacity. Second, the ecModel revealed a marked reduction in fluxes associated with ROS detoxification. This flux-level suppression is accompanied by the depletion of tocopheronic acid in rat metabolomics data, a metabolite linked to vitamin E–mediated antioxidant defense. Together, these findings provide a mechanistic explanation for the elevated oxidative stress observed in HSHF-fed livers, reflecting a dual burden of increased ROS production and impaired detoxification capacity. Third, under HSHF conditions, reduced fluxes were observed in leucine catabolism and estrogen metabolism subsystems, suggesting a diminished capacity for amino acid utilization and hepatic hormonal clearance under dietary stress. Collectively, these results indicate that HSHF drives a transition of hepatic metabolism from a flexible, multi-functional system toward a constrained, lipid-centric state characterized by mitochondrial stress, impaired redox control, and loss of systemic regulatory capacity.

While the ecModel-based analysis provides novel insights into tissue metabolic shifts, several limitations should be noted. First, as a genome-scale representation, the current model remains a simplified abstraction of *in vivo* metabolism; it may not fully capture post-translational modifications or allosteric regulation, both of which are critical for fine-tuning enzyme activities regardless of protein abundance. Furthermore, these models rely on steady-state assumptions. Integrating disease- or cell-type-specific ecGEMs with dynamic data in future studies may provide deeper insights into the progression of metabolic disorders.

## Acknowledgements

This work was financially supported by the Knut and Alice Wallenberg Foundation. The authors acknowledge the support from the Alpha Cell project.

## Contributions

Conceptualization: C.Z. and A.M. Investigation: M.L. and H.T. Formal analysis: M.L. and M.S. Writing – Original Draft: M.L. Supervision: A.M. and M.U. Writing – Review & Editing: All authors.

## Method

### Data collection and preprocessing

For tissue RNA-seq data, raw FASTQ files from 200 samples across 32 human tissues were downloaded from the EMBL-EBI database (accessions E-MTAB-2836 and E-MTAB-1733). Transcript abundance was quantified using Kallisto (v0.48.0)^11^ by pseudo-aligning reads to the GRCh38 human reference transcriptome. Gene-level expression estimates were then obtained by summarizing transcript abundances with tximport (v1.30.0)^12^. Single-cell RNA-seq data covering 82 cell types were downloaded from the Human Protein Atlas (HPA) single-cell resource. Normalized expression values (nTPM) for each cell type were used for downstream analyses^9^. Liver RNA-seq data from patients with metabolic-associated fatty liver disease (MAFLD) were downloaded from the GEO database (accession GSE135251)^13^.

### Human1 tissue specificity

To explore the tissue-specific features of enzyme-encoding genes in Human1, we integrated the Human1 gene list with the tissue-specific expression data from the HPA resource^7^. HPA classifies genes into three levels of tissue specificity: (i) tissue-enriched genes, defined as those with mRNA levels in one tissue at least five times higher than in all other analyzed tissues; (ii) group-enriched genes, which show elevated expression in a limited number of tissues; and (iii) tissue-enhanced genes, characterized by moderately increased expression across one or more tissues. By mapping Human1 genes to these HPA categories, we identified metabolic subsystems exhibiting high tissue specificity and further examined the tissue features of genes involved in multiple metabolic reactions.

In addition, Weighted Gene Co-expression Network Analysis (WGCNA) was performed on the expression profiles of 2,897 enzyme-encoding genes from Human1 using RNA-seq data from 200 samples representing 32 human tissues. A signed adjacency matrix was constructed based on pairwise gene correlations, with a soft-thresholding power of 12 applied to approximate a scale-free topology. Hierarchical clustering of the resulting topological overlap matrix (TOM) was then conducted, with the minimum module size set to 30. Genes within each identified module were functionally annotated using Gene Set Overrepresentation Analysis (GSOA) based on the Gene Ontology (GO) database to explore their biological relevance. To assess the tissue specificity of the co-expression modules, module eigengenes—representing the first principal component of each gene module—were correlated with tissue type indicators. Pearson correlation coefficients were calculated between module eigengenes and tissue indicator variables to quantify the association between each module and tissue type. Modules showing strong correlations (r > 0.7) were considered tissue-specific.

### ecGEM construction

Tissue-specific and cell-specific genome-scale metabolic models (GEMs) were constructed using the ftINIT algorithm based on Human1 (version 1.15). For each tissue or cell type, we calculated the mean gene expression value across multiple samples to generate a representative expression profile, which was then used to build the corresponding context-specific GEM. A gene expression threshold of 1 TPM was applied to define active reactions with gene–protein–reaction (GPR) associations. Reactions lacking GPRs were evaluated separately and retained only if necessary to maintain model feasibility (e.g., exchange or biomass reactions).

Enzyme-constrained genome-scale metabolic models (ecModels) were subsequently reconstructed following the GECKO 3 protocol^14^. The ecHuman-GEM model was built with a total protein pool constraint, and flux balance analysis (FBA) was performed to assess model feasibility. Finally, context-specific ecModels were generated by subsetting the generic ecHuman-GEM according to the corresponding tissue- or cell-specific GEMs.

For downstream analyses, we first compared the number of reactions identified in each tissue- and cell-specific ecGEM and visualized the results using line charts and heatmaps.t-SNE was applied to visualize structural differences among ecGEMs based on the presence or absence of reactions. To further investigate metabolic diversity across tissues and cell types, we identified the top 20 metabolic subsystems showing the highest variance in the number of reactions included across models.

### COMPASS infer metabolic activity

COMPASS was used to characterize metabolic activity across different tissues and cell types by integrating gene expression data. The expression levels of genes corresponding to each reaction were used to infer the relative activity of metabolic reactions^15^. The reference human metabolic model was derived from Recon3D, obtained from the BiGG Models platform^16^. Reactions associated with the Miscellaneous, Transport, and Extracellular Exchange subsystems were removed before downstream analysis.

Spearman correlations were calculated between each tissue–cell pair to assess similarities in metabolic activity based on the 6,135 identified metabolic reactions. The representative cell types showing the highest correlation with their corresponding tissues were then identified. A Kruskal–Wallis test was applied to evaluate differences in metabolic subsystem activity— including glycolysis, oxidative phosphorylation, fatty acid oxidation, and alanine and aspartate metabolism—across tissues. To further investigate tissue-specific metabolic differences, we used the limma package (version 3.58.1)^17^ to test for differential activity between each tissue and all remaining tissues, with p-values adjusted for multiple testing using the Benjamini–Hochberg (BH) method.

### Flux balance analysis of liver-ecGEM

Based on the developed liver-ecGEM, we performed a two-step flux balance analysis (FBA) to infer metabolic flux changes under two dietary conditions: a high-sugar and high-fat diet (HSFD) and a HSFD-constrained diet. Exchange reactions corresponding to major glucose and lipid-related carbon sources—including long-chain fatty acids (palmitate, oleate, stearate), short-chain organic acids (acetate), sterols (cholesterol), and lipid backbone molecules (glycerol)—were identified based on their reaction IDs in the model. For each condition, the lower bounds of the corresponding exchange reactions were adjusted to reflect the maximum allowed uptake rates. Each simulation was performed in two stages. In the maximal growth phase, biomass production was set as the objective function and maximized to determine the maximum growth rate. In the minimal protein usage phase, the biomass flux was constrained to 99% of its maximal value, and total protein usage (prot_pool_exchange) was minimized to estimate enzyme demand under near-optimal growth conditions. The resulting flux distributions were extracted and compared between the two dietary conditions to assess metabolic adjustments in response to nutrient availability.

### *In Vivo* Validation Using a Rat Dietary Model

To validate the metabolic changes inferred from the liver-ecGEM *in vivo*, a rat model was established under different dietary conditions. All animal procedures were approved by the Local Animal Experiments Ethics Committee (HADYEK) of Atatürk University (approval no. 2300106130). Female Sprague–Dawley rats (6–8 weeks old; 250 ± 17 g) were obtained from the Experimental Research and Application Center of Atatürk University. The control group (n = 8) received a standard laboratory diet (Bayramoğlu Yem, Erzurum, Turkey) throughout the study. The HSFD group (n = 7) was fed a high-sugar and high-fat diet (MD.88137, adjusted-calorie formulation, Supplementary Figure 3) to induce hypercholesterolemia. Liver fat accumulation was monitored by repeated magnetic resonance imaging (MRI) measurements. After 18 weeks of feeding, blood samples were collected from all animals for plasma metabolic analysis.

### Mass Spectrometric Detection and Data Acquisition

Plasma samples were processed for untargeted metabolomics and lipidomics analyses using high-resolution LC–MS.

For metabolomics, 50 µL of plasma was mixed with 400 µL of ice-cold acetonitrile/methanol/acetone (8:1:1, v/v/v) containing isotopically labeled internal standards (L-Valine-d8 and L-Phenyl-d5-alanine-2,3,3-d3; TRC Canada). After vortexing and incubation at 4 °C for 1 h, samples were centrifuged (16,000 × g, 15 min, 4 °C), the supernatant was evaporated under nitrogen, and reconstituted in LC–MS-grade water.

For lipidomics, 50 µL of plasma was extracted using the BUME method with 1-butanol/methanol (1:1, v/v) containing 5 mM ammonium formate and Avanti LipidoMIX internal standards. After vortexing, sonication (60 min, 20 °C), and centrifugation, the supernatant was dried under nitrogen and reconstituted in 1-butanol/methanol (1:1, v/v). Chromatographic separation was achieved on a Vanquish Horizon UHPLC system (Thermo Fisher Scientific) equipped with either a C18 Hypersil GOLD (metabolomics) or C30 Accucore (lipidomics) column, both maintained at 45 °C.

Mass spectrometric detection was performed using an Orbitrap Exploris 240 operated in positive and negative electrospray ionization modes (HESI). Data were acquired at 120,000 resolution (m/z 67–1700) using an AcquireX Deep Scan acquisition strategy with EASY-IC internal calibration.

Raw data were processed using Compound Discoverer 3.3 (Thermo Fisher Scientific) for peak detection, alignment, and compound identification against mzCloud, HMDB, ChemSpider, and Lipid MAPS databases. Lipid annotation was assisted by LipidSearch 4.2, and quality control correction was applied using SERRF normalization.

### LC–MS Data Processing and Statistical Analysis

For downstream LC–MS data analysis, normalized intensity values were used. Blank samples were applied for background correction by subtracting the mean signal of blanks from each feature, with negative values replaced by zero. Subsequently, quality control (QC) normalization was conducted by dividing each feature intensity by its mean value across all QC samples to minimize batch effects and instrumental drift. Features with a relative standard deviation (RSD) ≥ 40% in QC samples were considered unstable and removed. In total, 1,158 metabolite features and 2,581 lipid features were identified. Principal component analysis (PCA) was performed to visualize differences in metabolic profiles using the pcaMethods R package (version 1.94.0)^18^. Student’s t-test, followed by Benjamini–Hochberg (BH) correction, was applied to identify significantly different metabolites between groups.

### MAFLD cohort analysis

Differential expression analysis between MAFLD and healthy controls was performed using the DESeq2 R package (v1.36.0)^19^. Significantly differentially expressed genes (DEGs) were identified using an adjusted p-value threshold of < 0.05. The significantly upregulated and downregulated DEGs were further analyzed using Gene Set Overrepresentation Analysis (GSOA) based on the Gene Ontology (GO) database to identify significantly affected biological pathways.

